# *In vitro* prototyping and rapid optimization of biosynthetic enzymes for cellular design

**DOI:** 10.1101/685768

**Authors:** Ashty S. Karim, Quentin M. Dudley, Alex Juminaga, Yongbo Yuan, Samantha A. Crowe, Jacob T. Heggestad, Tanus Abdalla, William Grubbe, Blake Rasor, Dave Coar, Maria Torculas, Michael Krein, FungMin (Eric) Liew, Amy Quattlebaum, Rasmus O. Jensen, Jeff Stuart, Sean D. Simpson, Michael Köpke, Michael C. Jewett

## Abstract

Microbial cell factories offer an attractive approach for production of biobased products. Unfortunately, designing, building, and optimizing biosynthetic pathways remains a complex challenge, especially for industrially-relevant, non-model organisms. To address this challenge, we describe a platform for *in vitro* <u>P</u>rototyping and <u>R</u>apid <u>O</u>ptimization of <u>B</u>iosynthetic <u>E</u>nzymes (iPROBE). In iPROBE, cell lysates are enriched with biosynthetic enzymes by cell-free protein synthesis and then metabolic pathways are assembled in a mix-and-match fashion to assess pathway performance. We demonstrate iPROBE with two examples. First, we tested and ranked 54 different pathways for 3-hydroxybutyrate production, improving *in vivo* production in *Clostridium* by 20-fold to 14.63 ± 0.48 g/L and identifying a new biosynthetic route to *(S)*-(+)-1,3-butanediol. Second, we used iPROBE and data-driven design to optimize a 6-step *n*-butanol pathway, increasing titers 4-fold across 205 pathways, and showed strong correlation between cell-free and cellular performance. We expect iPROBE to accelerate design-build-test cycles for industrial biotechnology.

## Introduction

For decades, scientists and engineers have turned to biological systems to help meet societal needs in energy, medicine, and materials—especially when chemical synthesis is untenable (e.g., antimalarial drugs). Often, biologically-produced small molecules are insufficient for production at the optimal titer, rate, or yield because natural sources are difficult to optimize and may not scale easily (e.g., plants grow slowly). Thus, engineers seek to design enzymatic reaction schemes in model microorganisms to meet manufacturing criteria.^1^ Success in these endeavors depends upon identifying sets of enzymes that can convert readily available molecules (e.g., glucose) to high-value products (e.g., medicines), with each enzyme performing one of a series of chemical modifications. Unfortunately, this is difficult because design-build-test (DBT) cycles—iterations of re-engineering organisms to test new sets of enzymes—are detrimentally slow.^2,3^ As a result, a typical project today might only explore dozens of variants of an enzymatic reaction pathway. This is often insufficient to identify a commercially relevant solution because selecting productive enzymes using existing single-enzyme kinetic data has limited applicability in multi-enzyme pathways and consequently requires more DBT iterations.^4^ This challenge is exacerbated in industrially-relevant, non-model organisms (such as clostridia) for which genetic tools are not as sophisticated, high-throughput workflows are often lacking, there exist transformation idiosyncrasies, and there is reduced availability of validated genetic parts.

Yet, many industrial bioprocesses (e.g., synthesis of amino acids,^5^ organic acids,^6-8^ solvents^9,10^) rely on non-model organisms as they offer exceptional substrate and metabolite diversity, as well as tolerance to metabolic end-products and contaminants, making them excellent chassis for biochemical production of exotic molecules from an array of possible feedstocks. Clostridia specifically were used industrially in acetone-butanol-ethanol (ABE) fermentations in the early-to-mid 20^th^ century because of their unique solventogenic metabolism to produce large amounts of solvents (e.g., acetone and butanol) but were eventually phased out of use due to the success of petroleum until recently.^11^ In addition, cellulolytic clostridia can degrade lignocellulosic biomass and acetogenic clostridia can robustly ferment on a variety of abundant, low-cost C1 gases including waste gases from industrial sources (e.g., steel mills, processing plants or refineries) or syngas generated from any biomass resource (e.g., municipal solid waste or agricultural waste).^12^ Acetogenic clostridia have recently proven industrial relevance for full commercial scale ethanol production using emissions from the steel making process.^13^ However, these strains tend to lack the natural machinery to produce such solvents or other more complex products, and the tools to engineer them are underdeveloped with the solutions established for *E. coli* and yeast not being directly transferrable.^14^ In fact, until a few years ago, *Clostridium* organisms were considered genetically inaccessible with only a handful of genomic modifications being reported.^15,16^ While developing tools for engineering *Clostridium* is ongoing and promising progress has been made,^17,18^ discovering methods to speed up metabolic engineering DBT cycles for these organisms would accelerate the re-industrialization of such organisms.^14,19^

Cell-free systems provide many advantages for accelerating DBT cycles.^20-22^ For example, the open reaction environment allows direct monitoring and manipulation of the system to study pathway performance. As a result, many groups have used purified enzyme systems to study enzyme kinetics and inform cellular expression: testing enzymatic pathway performance *in vitro*, down-selecting promising pathway combinations, and implementing those in cells.^20,23-26^ Crude lysates are becoming an increasingly popular alternative to purified systems to build biosynthetic pathways because they inherently provide the context of native-like metabolic networks.^27-29^ For instance, the Panke group has shown that DHAP can be made in crude lysates and real-time monitoring can optimize production.^29^ In addition, our group has shown that 2,3-butanediol,^30^ mevalonate,^28^ *n-*butanol,^27,31^ limonene,^32,33^ and more complex products^34^ can be constructed in crude lysates with high productivities (>g/L/h). However, to our knowledge, no attempts have been made using cell-free prototyping to improve engineering of industrially-relevant, non-model organisms.

To address this opportunity, we report a new *in vitro* <u>P</u>rototyping and <u>R</u>apid <u>O</u>ptimization of <u>B</u>iosynthetic <u>E</u>nzymes approach (termed iPROBE) to inform cellular metabolic engineering. The foundational principle is that we can construct discrete enzymatic pathways through modular assembly of cell lysates containing enzymes produced by cell-free protein synthesis rather than by living organisms (**Figure 1**). This reduces the overall time to build pathways from weeks (or even months) to a few days, providing an increased capability to test numerous pathways by avoiding inherent limitations of cell growth and thus diminishing the reliance on single-enzyme kinetic data. A key conceptual innovation is that the DBT unit can be cellular lysates rather than genetic constructs, allowing us to perform DBT iterations without the need to re-engineer organisms. The rapid ability to build pathways *in vitro* using iPROBE allows us to generate large amounts of data describing pathway operation under several operating conditions. We demonstrate iPROBE in two ways. First, we use a new quantitative ranking system to bridge cell-free data and cellular metabolic engineering for the production of 3-hydroxybutyrate (3-HB) in *Clostridium autoethanogenum* from C1 gas. Specifically, we tested 54 different enzyme combinations for 3-HB and identify pathway combinations that produce at high-titers *in vivo*. The work also led to identification of a new route to *(S)*-(+)-1,3-butanediol (1,3-BDO), both non-native products for acetogenic clostridia and to our knowledge the first demonstration for biological production of the *(S)*-(+)-isomer of 1,3-butanediol. Second, we show the utility of iPROBE by increasing cell-free *n*-butanol production ∼4-fold in less than two weeks by assessing performance of 205 pathways using data-driven design-of-experiments. We then selected 9 pathway combinations from the iPROBE screen to assess butanol production in *C. autoethanogenum* strains, observing a strong correlation (r^2^ > 0.9) between in cell and cell-free pathway performance.

**Figure 1.**
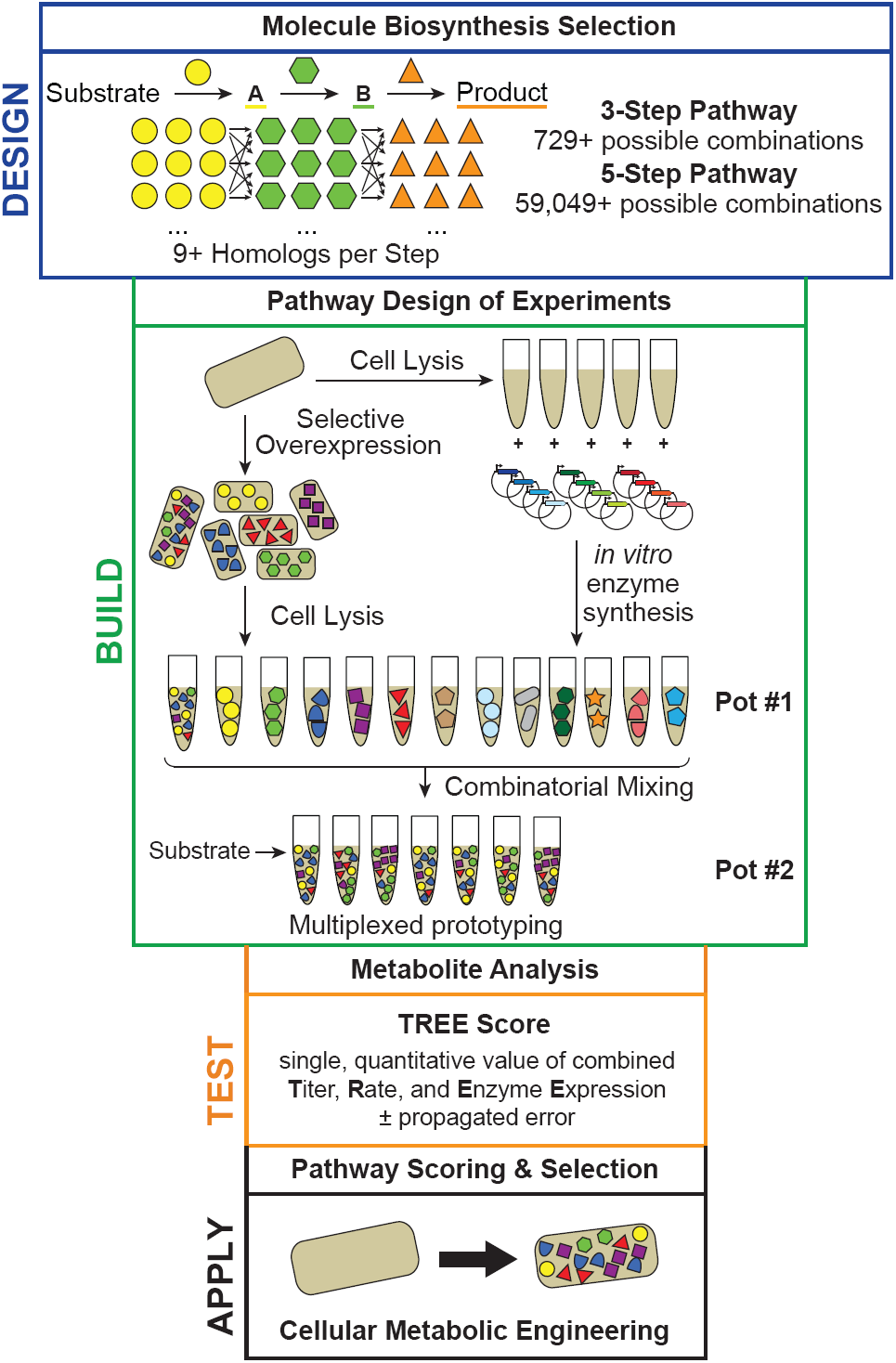
A two-pot cell-free framework for *in vitro* prototyping and rapid optimization of biosynthetic enzymes (iPROBE). A schematic overview of the iPROBE approach following a DBT and apply framework is depicted. In the design phase, reaction schemes and enzyme homologs are selected. In the build phase, lysates are enriched with pathway enzymes via overexpression prior to lysis or by cell-free protein synthesis post lysis. Then, lysates are mixed to assemble enzymatic pathway combinations of interest. In the test phase, metabolites are quantified over time and data is reduced into a single quantitative metric for pathway combination ranking and selection. In the apply phase, cell-free pathway combinations are selected and implemented in cellular hosts.

## Results

### Establishing a two-pot, cell-free framework for pathway prototyping

We selected 3-hydroxybutyrate (3-HB) biosynthesis as a prime example for pathway prototyping with iPROBE given it is non-native to clostridia and its importance as a high-value specialty chemical.^35,36^ Our vision was to demonstrate modular assembly of the pathway by mixing multiple crude cell lysates each individually enriched with a pathway enzyme, identify best sets of enzymes and their stoichiometry for pathway operation, and inform cellular design in an industrially proven,^12^ non-model host organism, in this case acetogenic *Clostridium autoethanogenum* (**Figure 1**). A unique feature of the iPROBE approach, relative to previous works in crude lysate-based cell-free prototyping,^27,31-33,37^ is that pathways are assembled in two steps (i.e., 2 pots), where the first step is enzyme synthesis via cell-free protein synthesis (CFPS) and the second step is enzyme utilization via substrate and cofactor addition to activate small molecule synthesis. Separating enzyme synthesis and utilization into two separate pots enables the modular control of enzyme concentrations because exact amounts can be added to the biosynthetic reaction. Moreover, negative physiochemical effects of the CFPS reaction^31^ on small molecule biosynthesis can be reduced.

We first set out to use iPROBE to study the impact of enzyme stoichiometry on pathway performance for the two, non-native enzyme pathway to 3-HB (**Figure 2A**). A thiolase (Thl) and a hydroxybutyryl-CoA dehydrogenase (Hbd), along with a native thioesterase,^38^ are required to make 3-HB. We initially selected a Thl gene from *Clostridium acetobutylicum* (*Cac*) and a Hbd gene from *Clostridium kluyveri* (*Ckl*) (**Supplementary Table S1**). We used the well-characterized *E. coli-*based PANOx-SP CFPS system^39^ to produce *Cac*Thl and *Ckl*Hbd, with soluble yields of 5.85 ± 0.82 µM and 19.31 ± 3.65 µM, respectively. Then, we designed five unique pathway combinations titrating different concentrations of Thl while maintaining a constant concentration of Hbd by mixing different ratios of CFPS reactions (keeping total CFPS reaction added constant using blank reactions containing no protein produced *in vitro*) (**Figure 2B-C**). Upon incubation with essential substrates, salts, and cofactors (e.g., glucose, NAD, CoA, ATP), we assessed 3-HB synthesis at 0, 3, 4, 5, 6, and 24 h for each of the five pathway combinations via high performance liquid chromatography (HPLC) (**Figure 2D**). The cell lysate contains endogenous enzymes for glycolysis that regenerate NADH^40^ and convert glucose to acetyl-CoA, providing the starting intermediate for 3-HB biosynthesis. As expected, no 3-HB was produced in the absence of Thl. The highest 3-HB titers were observed for 0.5 μM *Cac*Thl and 0.5 μM *Ckl*Hbd. We performed a similar titration of *Ckl*Hbd while maintaining a constant concentration of *Cac*Thl (**Supplementary Figure S1**). Taken together, our data show that crude lysates enriched by CFPS can be used to assemble metabolic reactions and sets the stage to optimize pathways using a two-pot cell-free approach.

**Figure 2.**
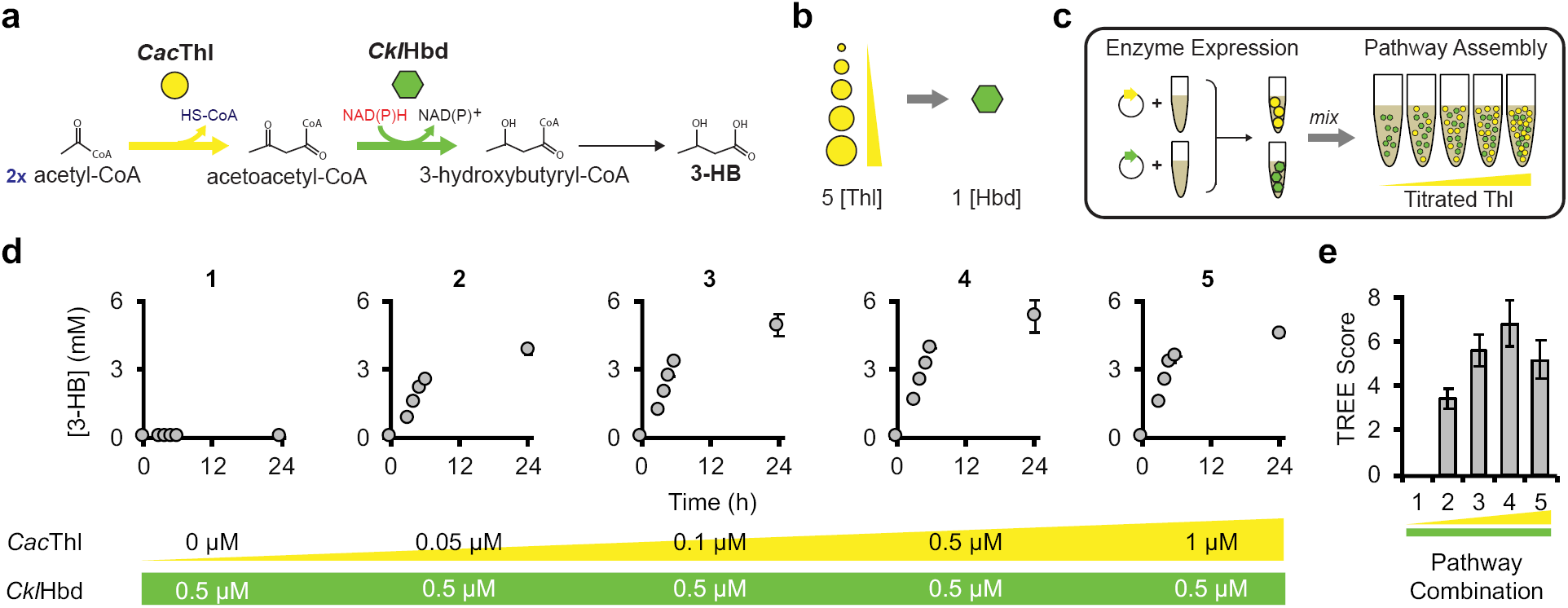
Individual pathway enzymes can be tuned in pathway context and ranked using TREE scores with iPROBE. (A) The pathway to produce 3-HB from native metabolism (acetyl-CoA) is selected requiring two enzymes not natively present, *Cac*Thl and *Ckl*Hbd. (B) Five pathway combinations are designed to be built and tested varying the concentration of *Cac*Thl low to high while maintaining *Ckl*Hbd at one concentration. (C) The five pathway designs are built by enriching two *E. coli* lysates with *Cac*Thl and *Ckl*Hbd, respectively, by CFPS (Pot #1). Then, the five pathway combinations are assembled by mixing CFPS reactions containing *Cac*Thl, *Ckl*Hbd, and no enzyme (blank) with fresh *E. coli* lysate. Kanamycin, to stop further protein synthesis, glucose and cofactors are added to start biosynthesis of 3-HB. (D) 3-HB is measured at 0, 3, 4, 5, 6, and 24 h after the addition of glucose for each of the five pathway combinations. Error bars are shown at 24 h and represent technical triplicates. (E) The TREE is score is then calculated for each pathway combination with propagated error.

### Developing a metric to quantify biosynthetic pathway performance

To optimize pathways with iPROBE, we next defined a pathway ranking system that would enable assessment of activity in the cell-free environment and hold potential to inform cellular design. The basis of this ranking system is a single, quantitative metric for our cell-free experiments. We call this metric the TREE score (for titer, rate, and <u>e</u>nzyme <u>e</u>xpression; important metrics for defining the success of cell-based metabolic engineering). The TREE score combines, through multiplication, titer at reaction completion, rate during the most productive phase of pathway operation, and enzyme expression as measured by protein solubility and total enzyme amount. Using our initial set of data (**Figure 2**) as a guide, the TREE score is obtained by multiplying 3-HB titer at 24 h, the linear 3-HB production rate between 3 and 6 h, and the sum of the average solubility of the pathway enzymes, Thl and Hbd, and the inverse of the total enzyme concentration for each of the five pathway combinations (**Supplementary Figure S2**; **Figure 2E**). While the TREE score rankings are not drastically different from the titers or rates alone, which they should not be, they exaggerate differences that might arise from each component of the score. Combining titer and rate enables use of both in ranking cell-free pathway performance which is helpful as it is unknown whether cell-free titer or rate is more or less important for informing cellular metabolic engineering. Incorporating enzyme expression allows pathways that have expression difficulties to decrease the overall pathway rank so as to avoid enzymes with poor expression properties. By reducing the complexity of available cell-free data to one value, the TREE score enables a rapid approach to rank pathways for iPROBE.

### iPROBE informs selection of genetic regulatory architectures in Clostridium

With a pathway ranking system at hand, we next aimed to validate that cell-free experiments could generate design parameters for DNA construction of biosynthetic pathways in cells. Gene expression involves designing a coding sequence, 5’ and 3’ regulatory elements, and vector maintenance components, among other parts. Selection of the promoter regulatory strengths (e.g., high, medium, low) for the expression of a coding sequence, in particular, is an essential factor for pathway tuning. Thus, we set out to develop a correlation between specific enzyme concentrations in iPROBE and specific strength regulatory architectures, relative promoter strengths and plasmid copy number for a single operon comprising the 3-HB pathway, for expression in *C. autoethanogenum*. To achieve this goal, we built cell-free pathway combinations for 3-HB by co-titrating seven different enzyme concentrations of Thl and Hbd in our reactions (**Supplementary Figure S3**). Specifically, we built seven cell-free reactions in increasing total concentration added, combining *Cac*Thl and *Ckl*Hbd at equimolar amounts. We ran each cell-free reaction for 24 h and measured the titer of 3-HB produced (**Supplementary Figure S3C**). We observed that as the amount of enzyme added increases, the amount of 3-HB increased up to a threshold amount of 1 µM of each enzyme added. In parallel, we constructed plasmids expressing *Cac*Thl and *Ckl*Hbd under eight regulatory architectures (change in promoter strength and plasmid copy number) of increasing strength and transformed them into separate strains of *C. autoethanogenum*. We ran small-scale bottle fermentations of each strain under anaerobic conditions on carbon monoxide (CO), hydrogen (H_2_), and carbon dioxide (CO_2_) gas and measured stationary phase titers of 3-HB (**Supplementary Figure S3D**) and observed a similar trend to the cell-free studies. Namely, we found that increases in expression strength led to higher 3-HB titers. These data allowed us to build a cell-free to cell correlation that connects cell-free 3-HB production and corresponding enzyme concentrations to cellular 3-HB production and corresponding plasmid regulatory strength. We found that using < 0.1 µM enzyme *in vitro* corresponds to low regulatory strengths *in vivo*, using 0.1 - 0.3 µM enzyme *in vitro* corresponds to medium strengths *in vivo*, and using > 0.3 µM enzyme *in vitro* corresponds to high strengths *in vivo*. In principle, this allows us to screen many different pathway combinations in cell-free systems and provide a rational recommendation for plasmid construction of those pathway combinations in *Clostridium*, which is what we did next.

### *iPROBE can inform the selection of pathway enzymes in* Clostridium

To showcase the iPROBE approach, we next screened several possible 3-HB pathway combinations using cell-free experiments, ranked a subset of candidate cellular pathway combinations using the TREE score, and showed cellular *C. autoethanogenum* 3-HB biosynthesis from CO/H_2_/CO_2_ gas correlates with cell-free experimental results. To do this, we tested six enzyme homologs of each Thl and Hbd originating from different *Clostridium* species (**Figure 3A**; **Supplementary Table S1**). We selected all pathway combinations of the 12 enzymes keeping a fixed total concentration of enzyme added (high expression levels) to build in cell-free reactions (**Figure 3B-C**). By measuring 3-HB production over the course of 24 h along with soluble enzyme expression for each of the enzymes (**Supplementary Figure S4**), we are able to calculate TREE scores for each of the 36 pathway combinations (**Figure 3D**). We found that a majority of pathway combinations perform poorly with only a handful achieving TREE scores above a value of 2. We wanted to use this information to down-select the number of combinations to test in acetogenic clostridia due to the current limitations in high-throughput strain testing. We selected a subset of four pathway combinations from the iPROBE screening to test in *C. autoethanogenum* labeled A, B, C, and D. We constructed and transformed DNA with strong regulatory architectures and each of the four-pathway enzyme sets into separate strains of *C. autoethanogenum* (**Figure 3E**). We ran small-scale bottle fermentations of each strain under anaerobic conditions on CO/H_2_/CO_2_ gas mixture and measured 3-HB titers at four time-points during the fermentation (**Figure 3F**). We observed that the best cell-free pathway combination as determined by TREE score (D) also performs the best of the four in *Clostridium* cells, achieving 33.33 ± 1.44 mM. The worst pathway combination in cell-free experiments (A) is also the worst performer in *C. autoethanogenum*. The other two pathway combinations (B, C), which were not statistically different, fall in the middle in cellular experiments. At a high-level, these data suggest that the cell-free system is useful for guiding pathway selection in cells. This might be especially true for down-selecting pathway combinations that should not be tested in cells (i.e., produce little to no product). In fact, the best pathway designs tested in two recent studies that explored autotrophic 3-HB production in acetogenic clostridia produced ∼4 mM and ∼1 mM 3-HB.^41,42^ Their pathways correspond to TREE scores of 1.06 ± 0.07 and 0.02 ± 0.00, respectively. Based on our iPROBE screening we would have suggested not testing these combinations *in vivo*. For context, our best pathway had a TREE score of 17.76 ± 3.08. In sum, iPROBE offers a framework to rapidly design, build, and test pathway combinations in cell-free experiments in a matter of days, bypassing DNA construction and transformation limitations, and to facilitate implementation of promising pathway combinations for engineering success in cells.

**Figure 3.**
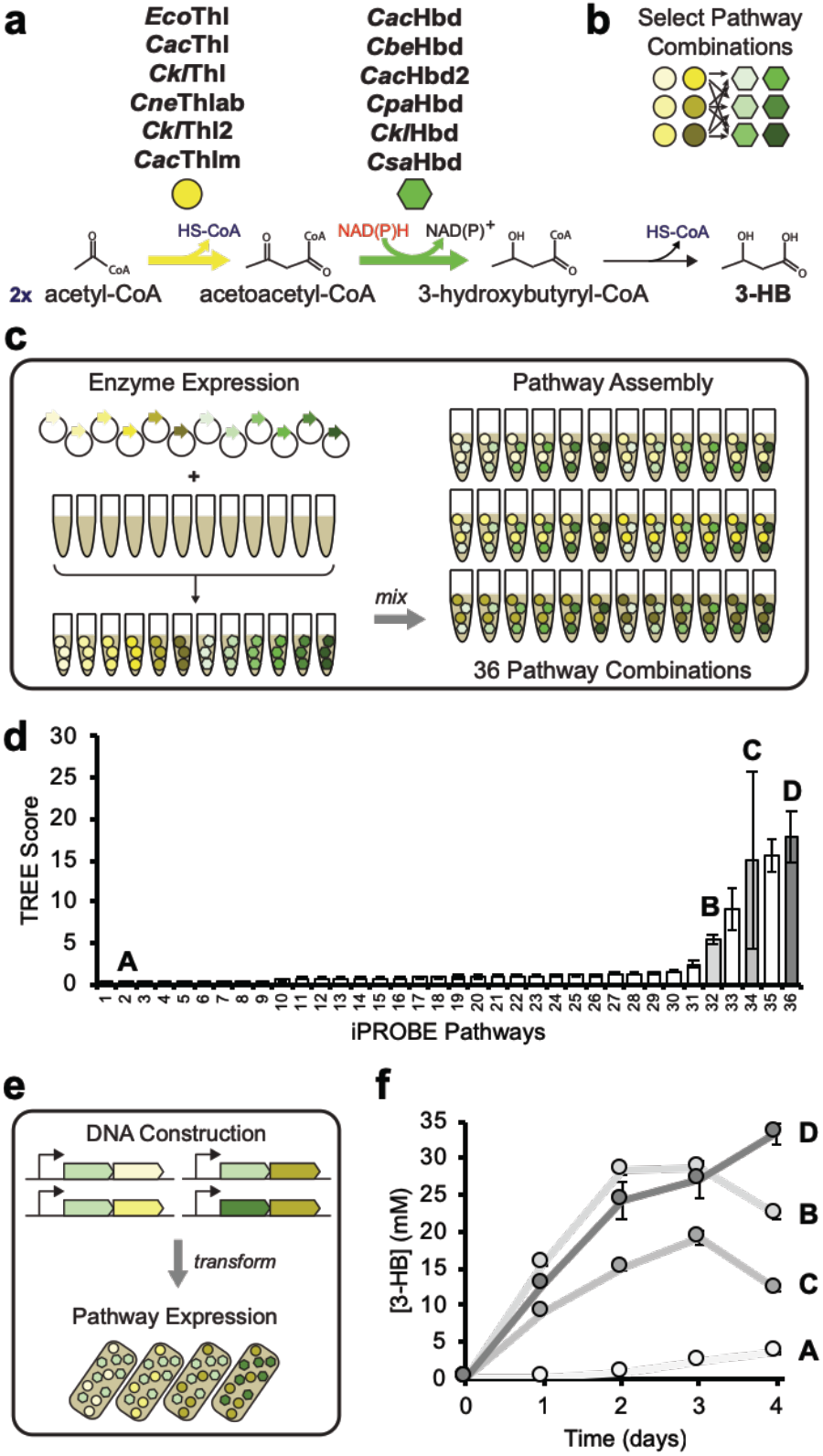
Enzymatic pathways can be screened with iPROBE to inform *Clostridium* expression for optimizing 3-hydroxybutyrate production. A reaction scheme for the production of 3-HB is presented in panel (a). Six homologs have been selected for each reaction step. The design in (b) includes the testing of six Thl homologs and six Hbd homologs at 0.5 µM each. We built each possible combination in cell-free systems (c) constituting 36 unique pathway combinations. We rapidly built these cell-free pathways by expressing each of the 12 enzyme variants in lysates by CFPS. We then mixed each to try all 36 possible combinations keeping enzyme concentration fixed. (d) 3-HB was measured, and TREE scores were calculated and plotted for each iPROBE pathway combination with propagated error. We then selected four pathway combinations to test in *C. autoethanogenum* (A, B, C, and D). These pathways were built in high copy plasmids with the highest strength promoters in single operons (e). *Clostridium* strains containing these pathway combinations were then fermented on gas and 3-HB was measured over the course of fermentation (f). Error bars represent technical triplicates.

### Scaled-up fermentations of 3-HB pathway prototype

We next coupled the results of our integrated iPROBE approach with in-house facilities to scale cell growth from bench to continuous fermentation scales (**Figure 4A**). Specifically, the best performing strain for 3-HB production selected by iPROBE was chosen for process scale-up from 0.1-L bottle fermentations to 1.5-L continuous fermentations using CO/H_2_/CO_2_ gas as the sole carbon and energy source. Over a 2-week fermentation, we monitored 3-HB and biomass in a control strain and our iPROBE-selected strain with and without optimized fermentation conditions (**Figure 4B-C**). In optimized fermentations we observed high-titers of 3-HB, ∼15 g/L (140 mM) at rates of >1.5 g/L/h in a continuous system. This is not only higher than the previously reported concentration in acetogenic *Clostridium*,^41,42^ but to our knowledge also exceeds the previously highest-reported concentration for traditional model organisms like *E. coli* (titer of ∼12 g/L and rate of ∼0.25 g/L/h in fed-batch)^38,43^ and yeast (titer of ∼12 g/L and rate of ∼0.05 g/L/h in fed-batch)^44^ without any additional genomic modifications to optimize flux into the pathway. This shows the utility of iPROBE in identifying pathways for industrial strain development. We anticipate that genome modifications to increase flux could further improve fermentation titers. For example, a recent study reported a 2.6-fold improvement in 3-HB production in a related engineered acetogenic *Clostridium* by downregulation of two native genes related to acetate production.^42^

**Figure 4.**
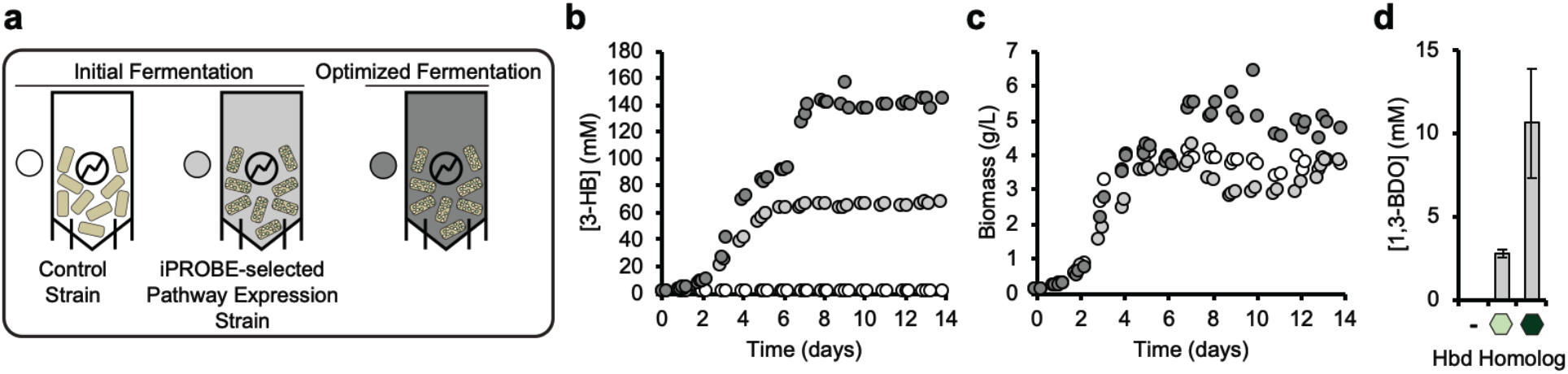
*Clostridium* fermentations show improved production of 3-HB and identification of a new route to 1,3-butanediol. (A) The iPROBE-selected optimal pathway, *Cac*Thl and *Ckl*Hbd1, for 3-HB production is built in a *C. autoethanogenum* strain and run in a 14-day continuous fermentation on CO/H_2_/CO_2_ gas as sole energy and carbon source. Fermentation parameter optimizations were performed for the new strain. Comparing fermentation of the control strain of *C. autoethanogenum* (white circles) to the initial (light grey circles) and optimized (dark grey circles) fermentations of the iPROBE-selected pathway expression strain, (B) 3-HB and (C) biomass were monitored. (D) 1,3-BDO was measured during steady-state fermentation and averages are shown for the control strain (-), the iPROBE-selected pathway expression strain (light green hexagon), and strain containing the PhaB homolog for Hbd (dark green hexagon).

Surprisingly, we also observed production of a novel metabolite, 1,3-butanediol (1,3-BDO), at 3-5% of the 3-HB levels and up to 0.5 g/L (**Figure 4D**). This is attributed to nonspecific activity of a native aldehyde:ferredoxin oxidoreductase (AOR) and alcohol dehydrogenase able to reduce 3-HB to 3-hydroxybutyraldehyde and further to 1,3-BDO. Indeed, no 1,3-BDO was observed when transforming the pathway into a previously generated AOR-knockout strain.^45^ These enzymes have previously been shown to reduce a range of carboxylic acids to their corresponding aldehydes and alcohols through reduced ferredoxin.^15,46^ While the *(R)*-(-)-form of 1,3-BDO has been produced via other routes,^47-49^ when using the *C. kluyveri*-derived Hbd we also detected the *(S)*-(+)-form of 1,3-BDO as determined by chiral analysis, which to our knowledge has never been produced in a biological system before. This chiral specificity is determined by the chosen 3-hydroxybutyryl-CoA dehydrogenase. Given that 1,3-BDO is used in cosmetics and can also be converted to 1,3-butadiene used in nylon and rubber production with a US$20 billion/year market,^35,50^ the discovery of this pathway is important. In sum, iPROBE provides a quick and powerful DBT framework to optimize and discover biosynthetic pathways for cellular metabolic engineering efforts, including those in non-model hosts.

### *Cell-free pathway prototyping for* n*-butanol biosynthesis*

Having demonstrated the use of iPROBE to optimize the 3-HB pathway for informing cellular design in *C. autoethanogenum*, we next aimed to show that iPROBE could be used to optimize longer pathways. We selected the 6-step pathway from acetyl-CoA to *n*butanol as a model pathway for this because butanol is an important solvent and drop-in fuel with US$5 billion/year market (**Figure 5A**). The key idea was to use iPROBE to optimize cell-free butanol production by constructing pathway variants with different enzyme homologs and reaction stoichiometry. The challenge with this optimization goal is the number of possible permutations. Indeed, testing just six homologs for each of the first four steps of the pathway at three different enzyme concentrations would alone require 314,928 pathway combinations, which exceeds typical HPLC analytical pipelines. Nevertheless, we know that homolog changes and reaction stoichiometry should be optimized simultaneously because tuning stoichiometries of one set of enzymes (*Eco*Thl, *Cbe*Hbd, *Cac*Crt, and *Tde*Ter; **Figure 5A**, highlighted in blue) does not achieve significant improvements (**Supplementary Figure S5**). To optimize enzyme homologs and concentrations simultaneously while also managing the landscape of testable hypotheses, we implemented a design-of-experiments using a neural-network-based algorithm to predict beneficial pathway combinations. This approach requires an initial data set to seed and train the neural networks.

**Figure 5.**
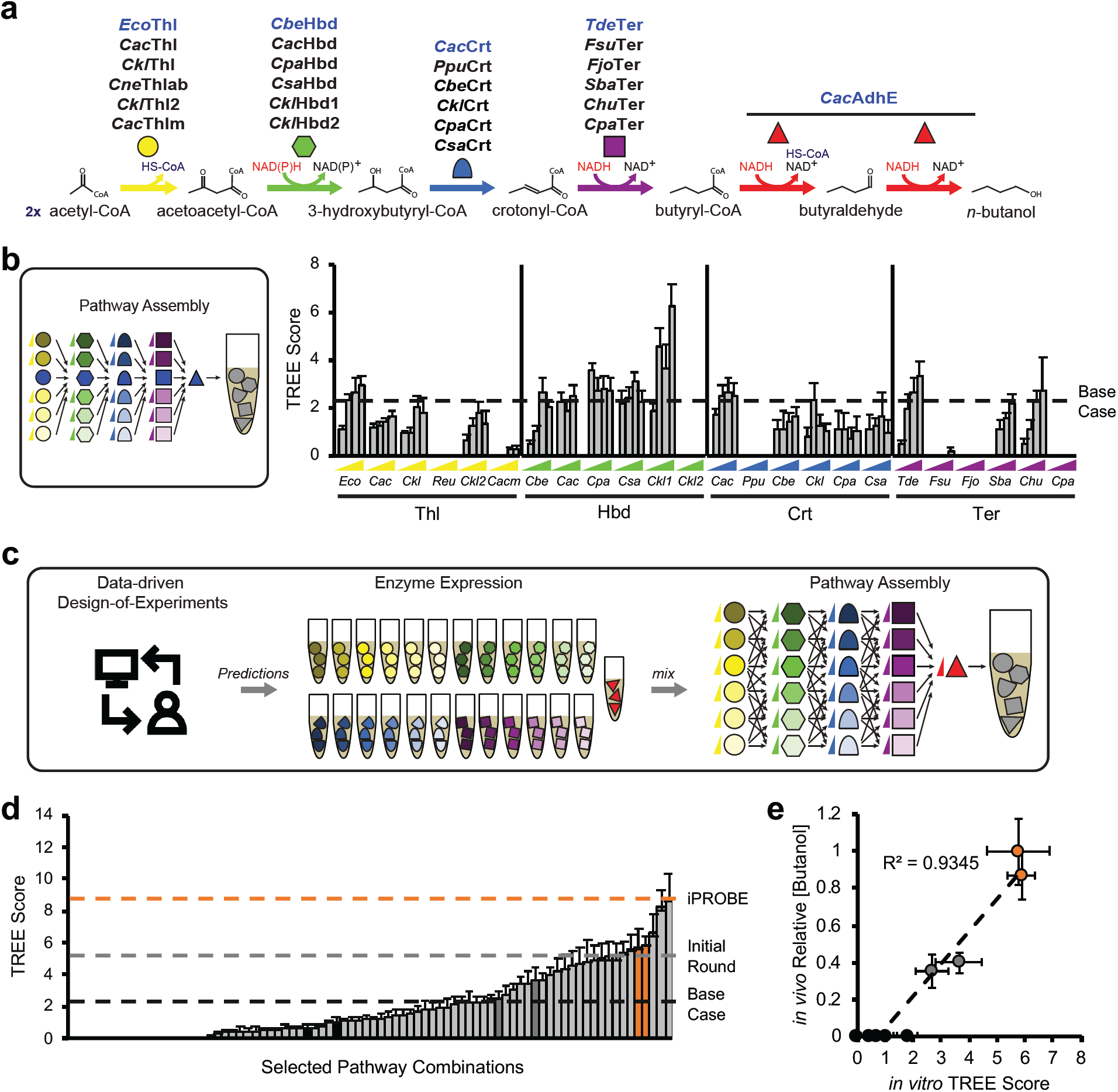
Cell-free pathway testing combined with data-driven design-of-experiments quickly screens 205 unique pathway combinations and selects pathways for cellular butanol production. A reaction scheme for the production of butanol is presented in panel (a). Six homologs have been selected for each of the first four reaction steps and are shown in panel (a). (b) The strategy for running an initial set of reactions is to test each homolog at 5 concentrations individually with the base case set of enzymes (blue). These 120 pathway combinations are run in cell-free reactions according to the two-pot iPROBE methodology and TREE scores are calculated from 24-h butanol time-courses. The dashed line is placed at the TREE score resulting from the base case set of enzymes. (c) Neural-network-based design of experiments is implemented using the data presented in (b) to predict enzyme sets and concentrations to be constructed. (d) These are then built in cell-free systems. TREE scores are calculated for all newly tested pathway combinations from 24-h butanol time-courses. The black dashed line is placed at the TREE score resulting from the base case set of enzymes, the grey dashed line corresponds to the TREE score of the best case from (b), and the orange dashed line represents the highest TREE score achieved through the data-driven iPROBE approach. (e) Nine pathway combinations (two high-performing pathways, two medium-performing pathways, and five low-performing pathways) were constructed and transformed *in vivo* in *C. autoethanogenum*. End-point titers of butanol production (relative to the highest *in vivo* titer) are plotted against *in vitro* TREE scores for the corresponding pathway combination. Error bars for *in vivo* titers are relative standard deviations for four to six replicates. Error bars for all TREE scores are propagated error based on TREE score calculations.

In creating the initial data set to guide improvements in titer, we chose six homologs of each Thl, Hbd, Crt, and Ter (**Supplementary Table S1**; **Figure 5A**). We tested five reaction stoichiometries for each enzyme homolog in a pathway context consisting of our base set of enzymes (**Figure 5A**, highlighted in blue), totaling 120 pathway combinations, including those in **Supplementary Figure S5**. We built these combinations in cell-free reactions from enzymes produced by CFPS, measured butanol production over time, and calculated TREE scores for each (**Figure 5B; Supplementary Figure S6**). In total, we collected these additional data in five experimental sets of 20 pathway combinations with each set taking five days (three days of HPLC time). A majority of the enzyme homologs did not out-perform the original enzyme set. This is not surprising as the base case has been extensively characterized and tested throughout the literature.^51-53^ However, we find that *Ckl*Hbd1 can double the TREE score at high concentrations and even out-performs the base case Hbd at lower concentrations. This agrees with an independent study that found a 1.6-fold improvement in ABE fermentation with *C. acetobutylicum* by replacing the native Hbd with the *Ckl*Hbd1.^54^

With this initial dataset collected (**Figure 5B**), we used ten randomly generated, trained, and evaluated neural network architectures to make pathway combination predictions (homolog set and reaction stoichiometry) to maximize TREE score which we could then build with the cell-free framework (**Figure 5C**). Design predictions suggesting enzyme concentrations of < 0.01 µM were ruled out, and we built the remaining 43 predictions in cell-free reactions (**Supplementary Figure S7A**). To evaluate our design-of-experiments approach, we compared the results with two additional sets of experiments: (i) a set varying reaction stoichiometry more thoroughly using only the base set enzymes (21 pathway combinations; **Supplementary Figure S7B**) and (ii) a hand-selected set of 18 pathway combinations based on our understanding of the biosynthesis (**Supplementary Figure S7C**). In total we tested 205 unique pathway combinations (base set combinations, initial round combinations, and data-driven designs) (**Figure 5D**; **Supplementary Figure S6**). Nearly 19% of the total pathway combinations screened have higher TREE scores than our base case. Without testing every possible combination, we were able to rapidly test a manageable subset achieving ∼4 times higher TREE scores (∼2.5 times higher titer and 58% increase in rate) over the base case pathway combination. The top six TREE scores each arose from pathways predicted in the neural-network-based design, showcasing the importance of merging computational and experimental design. Further analysis of the pathway combinations shows that there are enzyme homologs and concentrations that clearly improve the TREE score. The chosen thiolase does not seem to matter in the 20 top-performing combinations whereas the *C. kluyveri* Hbd1 is far superior to other Hbd enzymes (**Supplementary Figure S8A**). We also noticed that in the 20 top-performing combinations Hbd is present at significantly (p<0.001) higher concentrations and the median Crt concentration is lower, though not significantly, than the initial 0.3 µM (**Supplementary Figure S8B**). This suggests that optimal pathway operation occurs when enough flux can pass away from central carbon metabolism down the butanol pathway but minimal flux flows through the toxic intermediate metabolite, crotonyl-CoA.

We next assessed iPROBE’s ability to enhance cellular design by selecting nine representative pathway combinations from the iPROBE screening and constructing them in *C. autoethanogenum* strains to produce butanol: two pathway combinations scoring among the top five combinations, two pathway combinations in the middle range of the data set, and five pathway combinations near the tail-end of all combinations tested. To avoid diverting flux toward 3-HB, we identified and knocked-out a native thioesterase able to hydrolyze 3-HB-CoA from our screening strain. After monitoring butanol production over the course of six days (**Supplementary Figure S9A**), we see a high correlation (r^2^ = 0.93) between *in vivo* expression in *C. autoethanogenum* and TREE scores from iPROBE (**Figure 5E**). This further emphasizes that selecting top-performing pathways from iPROBE can improve production in *Clostridium* organisms and decreases the number of strains that need to be tested. While overall butanol production was low in *C. autoethanogenum* using the iPROBE-selected pathways, we were able to increase production over 200-fold from 0.1 ± 0.0 mM to 22.0 ± 1.3 mM (**Supplementary Figure S9B**) by replacing the trans-2-enoyl-CoA reductase (Ter) enzyme with the ferredoxin-dependent electron bifurcating enzyme complex (Bcd-EtfA:EtfB) naturally used for these activities in clostridia.^55^ This is not surprising in light of a recent study that showed Ter is detrimental to ABE fermentation when introduced in *C. acetobutylicum*.^56^ For comparison, the previously best reported butanol production in engineered acetogenic clostridia was ∼2 mM.^15^ Moreover, the Bcd-EtfA:EtfB complex is a delicate complex that is extremely oxygen sensitive^57^ and has so far been inactive in *E. coli* lysates (in alignment with previous reports that highlighted difficulties expressing Bcd in *E. coli*),^52^ highlighting an area for potential improvement of iPROBE (i.e., compatibility of *E. coli* lysates with non-model organisms). Overall, this work demonstrates the power of coupling data-driven design-of-experiments with a cell-free prototyping framework to select feasible subsets of pathways worth testing *in vivo* for non-model organisms.

## Conclusion

We demonstrate a new, two-pot framework, iPROBE, that incorporates a highly controllable and modular cell-free platform for constructing biosynthetic pathways with a quantitative metric for pathway performance selection (the TREE score) to engineer and improve small molecule biosynthesis in non-model organisms that can be arduous to manipulate. Specifically, we show that by screening 54 biosynthetic pathway combinations for the production of 3-HB in cell-free reactions, we can rationally select a handful of pathways to inform cellular metabolic engineering in clostridia. Indeed, iPROBE enabled the construction of a strain of *C. autoethanogenum* that produces high-titers and yields of 3-HB (∼20x higher than the previous highest reported concentration in the literature) in continuous fermentations with carbon monoxide/hydrogen/carbon dioxide gas as sole source of carbon and energy. The work also led to the identification of a new route to 1,3-BDO and the first production to our knowledge of the *(S)*-isomer of this molecule in a biological system. Beyond the 3-HB example, we also show the ability to use iPROBE in conjunction with data-driven design-of-experiments to reduce an exceedingly large landscape of testable pathway designs, test a subset of 205 pathway combinations *in vitro* for the production of butanol, and show that by testing a further subset of designs *in vivo* we can improve butanol production in acetogenic clostridia. Highlighting the utility of iPROBE for accelerating DBT cycles, the 205 pathway combinations for butanol were built cumulatively in 12 days (excluding HPLC time) in cell-free reactions, which we estimate would have taken a team of researchers more than 3 months in clostridia. Importantly, we show that there is indeed a correlation between pathway performance *in vitro* and *in vivo* providing evidence of the effectiveness of the iPROBE approach.

Looking forward, we anticipate that iPROBE will facilitate DBT cycles of biosynthetic pathways by enabling the rapid study of pathway enzyme ratios, tuning individual enzymes in the context of a multi-step pathway, screening enzyme variants for high-performance enzymes, and discovering enzyme functionalities. This in turn will decrease the number of the strains that need to be engineered and time required to achieve desired process objectives. This will increase the flexibility of biological processes to adapt to new markets, expand the range of fossil-derived products that can be displaced with bio-derived alternatives, and enhance the economic benefits for co-produced fuels.

## Methods

### Bacterial strains and plasmids

*Escherichia coli* BL21(DE3) (NEB) was used for preparation of cell extracts which were used to express all exogenous proteins *in vitro*.^37^ A derivate of *Clostridium autoethanogenum* DSM10061 obtained from the German Collection of Microorganisms and Cell Cultures GmbH (DSMZ; Braunschweig, Germany) was used for *in vivo* characterization and fermentations.^58^ For butanol production, this strain was used with a native thioesterase (CAETHG_1524) knockout made using Triple Cross recombination as described previously.^59^

Twenty-three enzymes were examined in this study (**Supplementary Table S1**). DNA for all enzyme homologs tested were codon adapted for *E. coli* using IDT codon optimizer. Non-clostridial sequences were codon adapted for *C. autoethanogenum* using a LanzaTech in-house codon optimizer, and all native clostridial genes were used as is. *E. coli* and *C. autoethanogenum* adapted sequences are listed in **Supplementary Table S2** and **Supplementary Table S3**, respectively. For the cell-free work, the pJL1 plasmid (Addgene #69496) was used. The modular pMTL80000 plasmid system^60^ along with *acsA*^45^, *fdx*^45^, *pta*^61^ and *pfor*^62^ promoters were used for the *C. autoethanogenum* plasmid expression.

### Cell Extract Preparation

*E. coli* BL21(DE3) cells were grown, harvested, lysed, and prepared using previously described methods.^27,63^

### iPROBE Reactions

Cell-free protein synthesis (CFPS) reactions were performed to express each enzyme individually using a modified PANOx-SP system described in previous pubications.^39,64^ Fifty-μL CFPS reactions were carried out for each individual enzyme in 2-mL microcentrifuge tubes. Enzyme concentrations in CFPS reactions were quantified by ^14^C-leucine incorporation during *in vitro* translation. Then reactions performed for identical enzymes were pooled together when multiple reaction tube-volumes were needed to keep a consistent 50-µL reaction volume and geometry for every CFPS reaction. Based on molar quantities of exogenous enzymes in each CFPS reaction determined by radioactive measurement, CFPS reactions were mixed to assemble complete biosynthetic pathways in 1.5-mL microcentrifuge tubes. CFPS reactions constitute 15 µL of a 30-µL-total second reaction. When the total CFPS reaction mixture constituted less than 15 µL, ‘blank’ CFPS reaction was added to make the total amount of CFPS reaction up to 15 µL. The ‘blank’ reactions consist of a typical CFPS reaction with no DNA added. This 15 µL CFPS mixture was then added to fresh extract (8 mg/mL), kanamycin (50 μg/ml), glucose (120 mM), magnesium glutamate (8 mM), ammonium glutamate (10 mM), potassium glutamate (134 mM), glucose (200 mM), Bis Tris pH 7.8 (100 mM), NAD (3 mM), and CoA (3 mM); final reaction concentrations are listed. Reactions proceeded over 24 h at 30 °C. Measurements from samples were taken at 0, 3, 4, 5, 6, and 24 h.

### Quantification of protein produced *in vitro*

CFPS reactions were performed with radioactive ^14^C-Leucine (10 µM) supplemented in addition to all 20 standard amino acids. We used trichloroacetic acid (TCA) to precipitate radioactive protein samples. Radioactive counts from TCA-precipitated samples was measured by liquid scintillation to then quantify soluble and total yields of each protein produced as previously reported (MicroBeta2; PerkinElmer).^39,40^

### Metabolite Quantification

High-performance liquid chromatography (HPLC) was used to analyze 3-HB and *n*-butanol. We used an Agilent 1260 series HPLC system (Agilent, Santa Clara, CA) via a refractive index (RI) detector. 3-HB and *n-*butanol were separated with 5 mM sulfuric acid as the mobile phase and one of two column conditions: (1) an Aminex HPX-87H or Fast Acids anion exchange columns (Bio-Rad Laboratories) at 35 or 55 °C and a flow rate of 0.6 ml min^-1^ or (2) a Alltech IOA-2000 column (Hichrom Ltd, Reading, UK) at 35 or 65 °C and flow rate of 0.7 ml min^-1^ as described earlier.^65^ 1,3-butanediol was measured using gas chromatography (GC) analysis, employing an Agilent 6890N GC equipped a Agilent CP-SIL 5CB-MS (50 m×0.25 μm×0.25 μm) column, autosampler and a flame ionization detector (FID) as described elsewhere.^65^ For chiral analysis of *(S)*-(+)-1,3-Butanediol and (R)-(-)-1,3-Butanediol an Agilent 6890N GC equipped with a Restek Rt®-bDEXse 30m x 0.25 mm ID x 0.25µm df column and a flame ionization detector (FID) was used. Samples were prepared by heating for 5 minutes at 80 °C, followed by a 3-minute centrifugation at 14,000 rpm. Exactly 400 µL of supernatant was then transferred to a 2-mL glass autosampler vial and 100 µL of and Internal Standard solution (5-methyl-1-hexanol and tetrahydrofuran in ethanol) was added. The capped vial was then briefly vortexed. Sample vials then were transferred to an autosampler for analysis using a 1.0 µL injection, a split ratio of 60 to 1, and an inlet temperature of 230 °C. Chromatography was performed with an oven program of 50 °C with a 0.5 min hold to a ramp of 3 °C/min to 70 °C to a ramp of 2 °C/min to 100 °C with a final ramp at 15 °C/min to 220 °C with a final 2-min hold. The column flow rate was 30 cm/sec using helium as the carrier gas. The FID was kept at 230 °C. Quantitation was performed using a linear internal standard calibration.

### TREE Score Calculations

The TREE Score is calculated by multiplying the titer by the rate by enzyme expression metric.

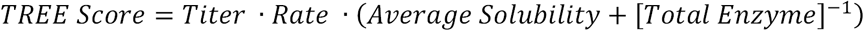

The titer is the metabolite concentration in the cell-free reaction at 24 h, when the reaction is complete. The error associated with titer is the standard deviation of reaction triplicates. The rate is the slope of the linear regression of metabolite concentrations taken at 3, 4, 5, and 6 h time points. The rate-associated error is the standard error of the slope calculated by the linear regression. The average solubility term is calculated by first determining the solubility (soluble protein/total protein, n = 3) for each individual enzyme via ^14^C-leucine incorporation. The average solubility is then the average value of enzyme solubilities (in this case, five enzymes) and the error associated with solubility is propagated error. The concentration of total enzyme is calculated by the addition of the final concentrations of each enzyme in the reaction with propagated error. The final error on the TREE score is the propagated error of each individual component.

### *In vivo* Gas Fermentations

*In vivo* cultivation and small-scale bottle fermentation studies were carried out as described earlier using a synthetic gas blend consisting of 50% CO, 10% H_2_, 40% CO_2_ (Airgas, Radnor, PA).^62^ Continuous fermentations were carried out in 1.5L continuous stirred tank reactors (CSTRs) with constant gas flow as described elsewhere.^65,66^

### Design-of-experiments using neural networks

A neural network-based approach was used to explore the vast landscape of possible experimental designs. The neural networks, which were all designed as deep neural network regressions, had between 5 and 15 layers and between 5 and 15 nodes in each layer. We randomly generated the network architectures, which we then evaluated based on how accurately the trained models predicted the training data. We trained and evaluated 500 unique network architectures and selected the top 10. Modeling enzymatic pathways requires a mix of continuous and categorical variables. Because many machine learning algorithms require numeric input and output variables, we used one-hot encoding, which is a process that converts categorical variables into a numerical form that machine learning algorithms can use. This method treats categorical variables as a multidimensional binary input that must sum to one. The concentration values were used as is, resulting in a 30 variable input matrix: 25 variables representing the categorical variation (i.e., different homologs) and 5 representing the concentration. Ten predictions were selected from each of the top 10 architectures and then we removed predictions that were impossible experimentally (i.e., concentrations too low to pipet accurate volumes).

### Statistics

All error bars on metabolite and protein quantification represent one standard deviation derived from technical triplicates. All error bars on TREE score values are propagated error as described in the TREE score calculation. In comparing the significance of enzyme concentration on TREE scores for butanol production in **Supplementary Figure 8B** we used the Mann-Whitney test to determine whether enzyme concentrations of the enzyme combinations that produced the top 20 TREE scores are greater than the enzyme concentrations of the entire data set.

### Data and Materials Availability

All cell-free data generated and shown in this manuscript are provided in **Supplementary File A** (.xlsx). Any additional data or unique materials presented in the manuscript may be available from the authors upon reasonable request and through a materials transfer agreement.

## Acknowledgments

We would like to thank Alexander P. Mueller, Ryan C. Tappel, Shivani Garg, Wyatt Allen, Loan Tran, and Steven D. Brown from LanzaTech for conversations regarding this work. In addition, we would like to thank Colleen Reynolds from Lockheed Martin for conversations on the design-of-experiments using neural networks. This work is supported by the U.S. Department of Energy, Office of Biological and Environmental Research in the DOE Office of Science under Award Number DE-SC0018249. We also thank the following investors in LanzaTech’s technology: BASF, CICC Growth Capital Fund I, CITIC Capital, Indian Oil Company, K1W1, Khosla Ventures, the Malaysian Life Sciences, Capital Fund, L. P., Mitsui, the New Zealand Superannuation Fund, Petronas Technology Ventures, Primetals, Qiming Venture Partners, Softbank China, and Suncor.

## Author contributions

ASK, SDS, MK, and MCJ designed the study; ASK, QMD, and MCJ developed the cell-free framework; ASK, SAC, JTH, WG, and BR performed all cell-free experiments; ASK and QMD analyzed cell-free data; AJ performed the *Clostridium* strain engineering for 3-HB and 1,3-BDO; TA performed the *C. autoethanogenum* gas fermentation for 3-HB and 1,3-BDO; YY, EL, ROJ and MK performed the *C. autoethanogenum* strain engineering and gas fermentation for Butanol; AJ, YY and MK analyzed *C. autoethanogenum* data; AQ developed analytical methods for 3-HB, 1,3-BDO, and Butanol; DC, MT, MKr, and JS performed all design-of-experiments using neural networks; ASK, MK, and MCJ wrote the manuscript.

## Supplementary Information

**Supplementary Figure S1.**
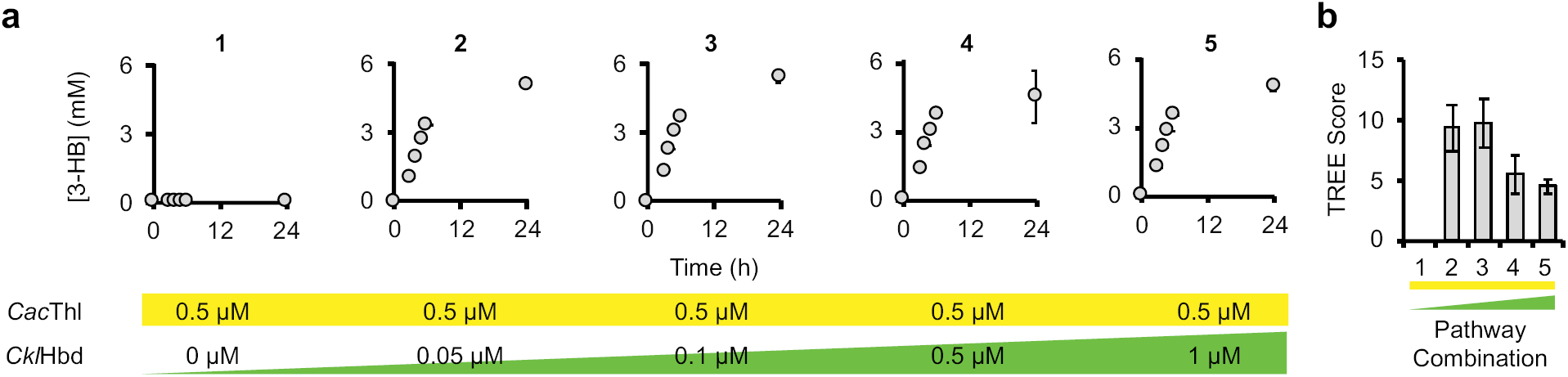
Hbd can be tuned in pathway context and assessed using TREE scores with iPROBE. Five pathway combinations are designed, built, and tested varying the concentration of CklHbd low to high while maintaining CacThl at one concentration. (A) 3-hydroxybutyrate is measured at 0, 3, 4, 5, 6, and 24 h after the addition of glucose for each of the five pathway combinations. Error bars are shown at 24 h and represent technical triplicates. (B) The TREE is score is then calculated for each pathway combination.

**Supplementary Figure S2.**
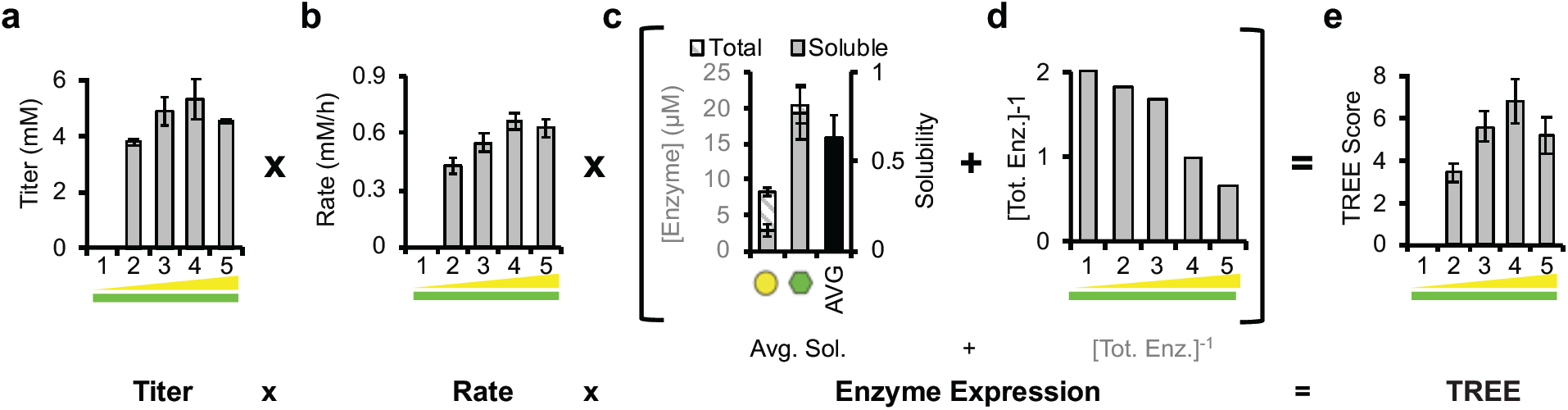
The titer, rate, and enzyme expression (TREE) score is calculated through multiplication. 3-hydroxybutyrate is measured at 0, 3, 4, 5, 6, and 24 h after the addition of glucose for each of the five pathway combinations described in **Figure 2**. From these measurements, 3-HB titer at 24 h (A) and rate of production through 6 h (B) is quantified. Error bars shown for titer represent technical triplicates. Error bars shown for rate represent the standard error of the linear regression. Enzyme expression is quantified by adding the average solubility of each enzyme (C) to the inverse of the total concentration of exogenous enzyme present (D). Error bars shown for enzyme concentrations represent technical triplicates. (E) The TREE is score is then calculated by multiplying the titer by the rate by the enzyme expression for each pathway combination with error bars representing the propagated error.

**Supplementary Figure S3.**
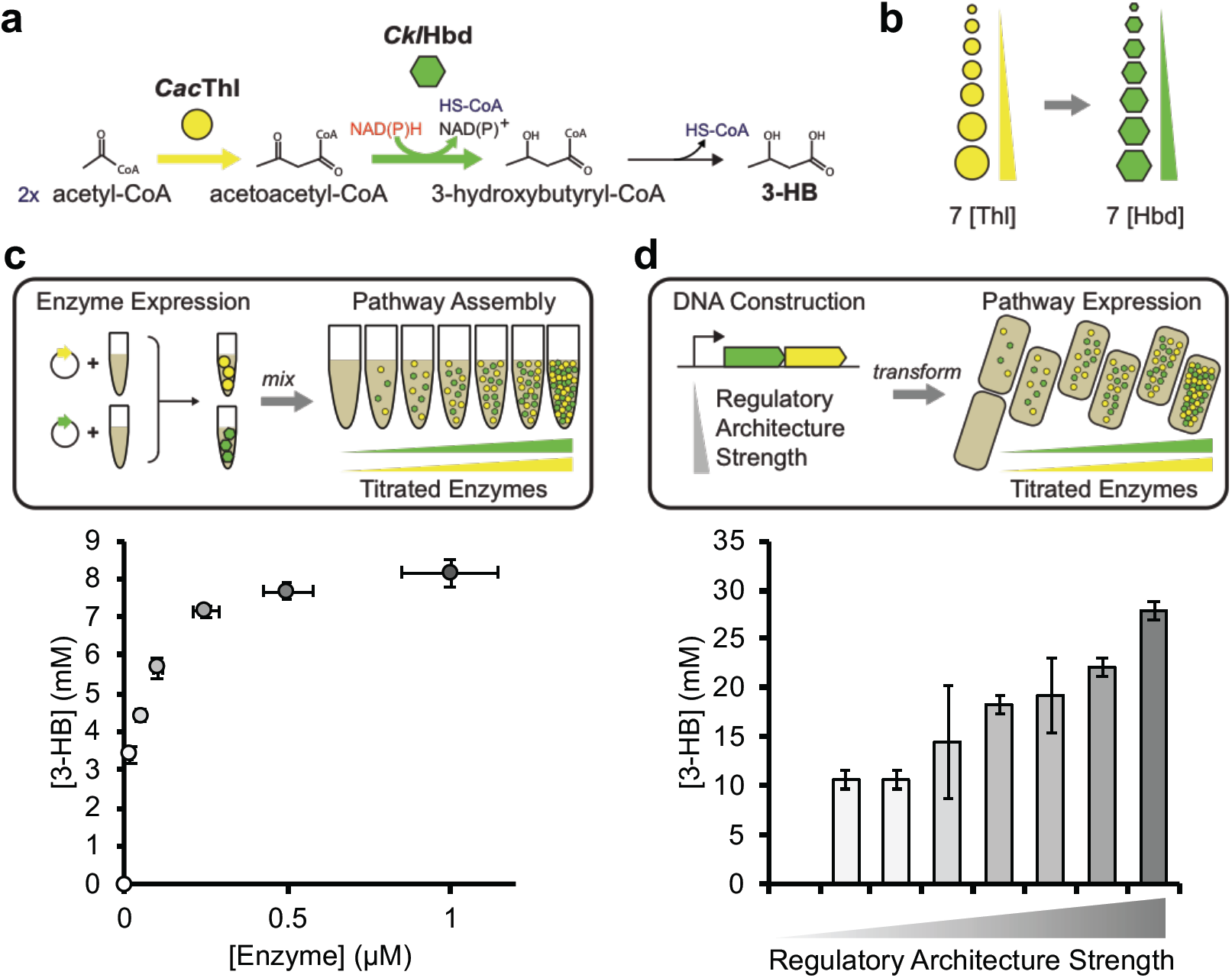
Enzyme concentrations can be tuned with iPROBE to inform genetic design for Clostridium expression of 3-hydroxybutyrate. A reaction scheme for the production of 3-HB is presented in panel (a). The design in (b) includes the co-titration of CacThl and CklHbd at seven concentrations (0, 0.02, 0.05, 0.1, 0.25, 0.5, and 1 µM). We built these seven designs in cell-free systems (c) by CFPS of each enzyme in separate lysates (Pot #1) followed by mixing to assemble full pathways for 3-HB production (Pot #2). We measured 3-HB over the course of 24 h for each. We compared these results to Clostridium-based expression by building eight genetic constructs with varying promoters and plasmid copy number (e). We measured final titer of 3-HB for each. Error bars represent technical triplicates. Error bars on enzyme concentrations are technical replicates with n > 3.

**Supplementary Figure S4.**
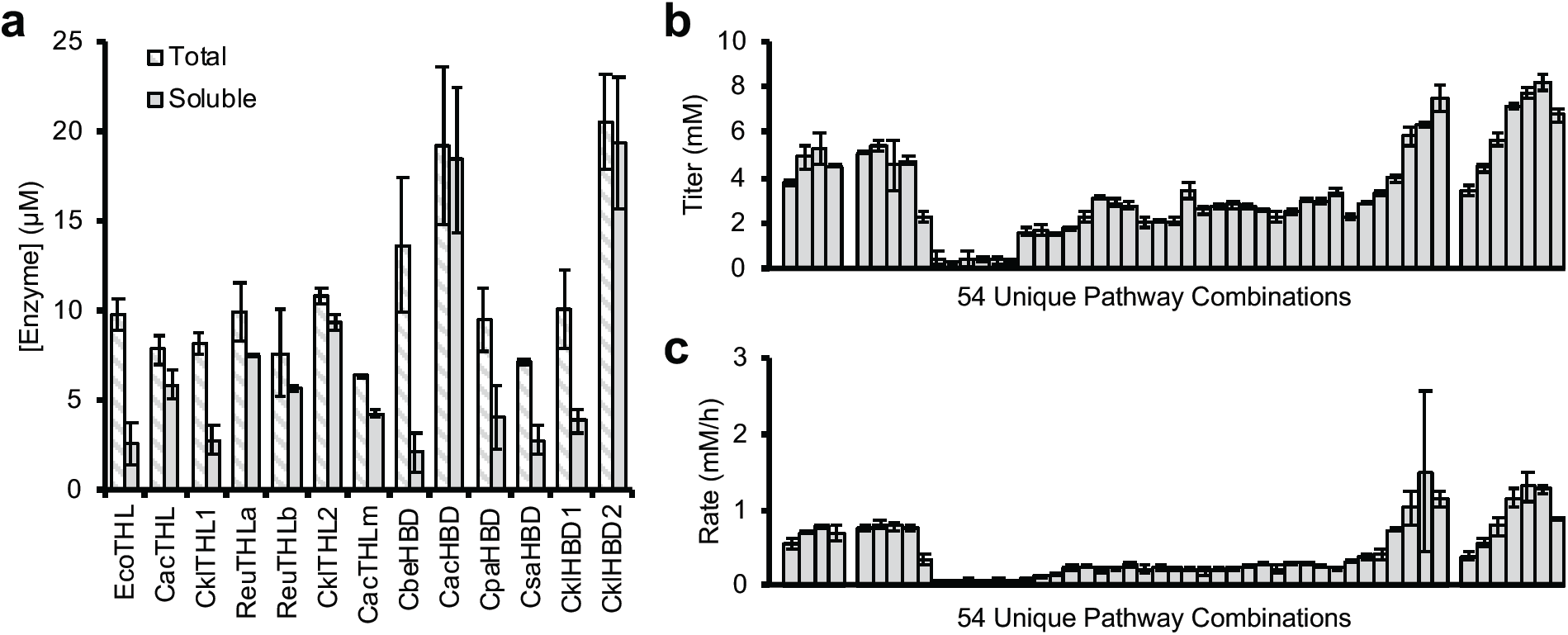
Cell-free 3-HB titers, rates, and enzyme expression. Six homologs have been selected for each reaction step (Thl and Hbd) for 3-HB production and expressed by cell-free protein synthesis. Total and soluble yields of protein are displayed in panel (a). Error bars represent technical triplicates. iPROBE was run in 54 combinations listed in Supplemental File A. Titers at 24 h are shown in panel (b) with error bars representing technical triplicates. Rates determined by linear regression between 3 h and 6 h measurements are shown in panel (c) with error bars representing the standard error of the linear regression.

**Supplementary Figure S5.**
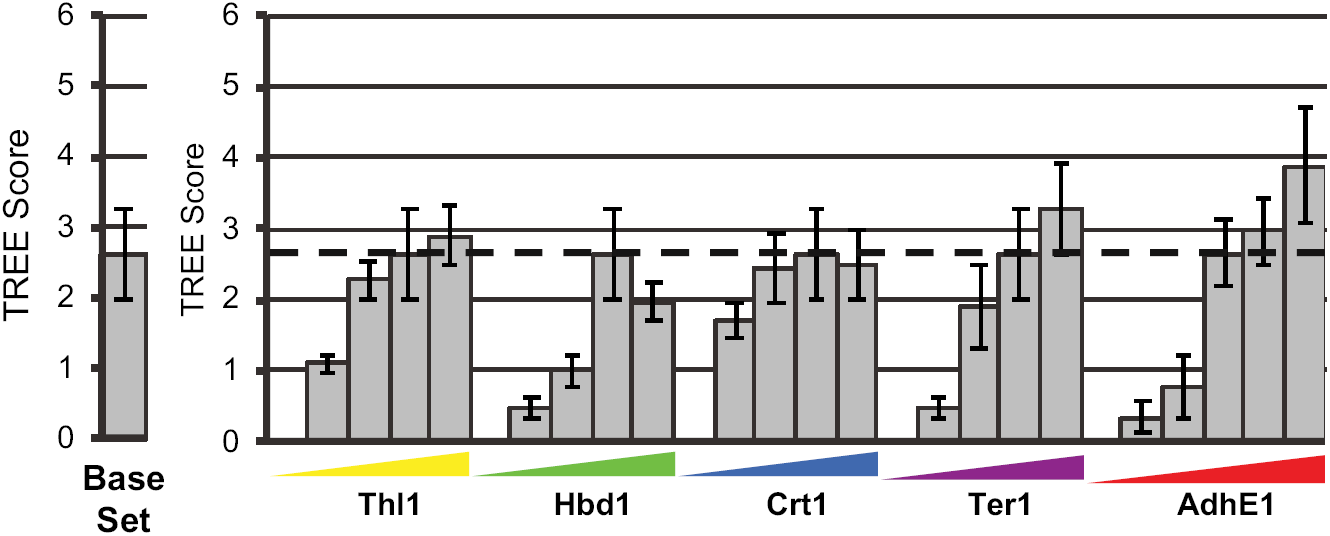
Titrating individual base case enzymes for the production of butanol. Cell-free reactions were run with concentrations of 0.3 µM for EcoThl, CbeHbd, CacCrt, and TdeTer while 0.6 µM was used for CacAdhE for the ‘Base Set’. Each enzyme concentration was altered individually to be 0, 0.1, 0.3, and 0.5 µM with the remaining enzymes remaining at the ‘Base Set’ values (essentially a titration of each enzyme). TREE scores are calculated for each combination with error bars representing propagated error.

**Supplementary Figure S6.**
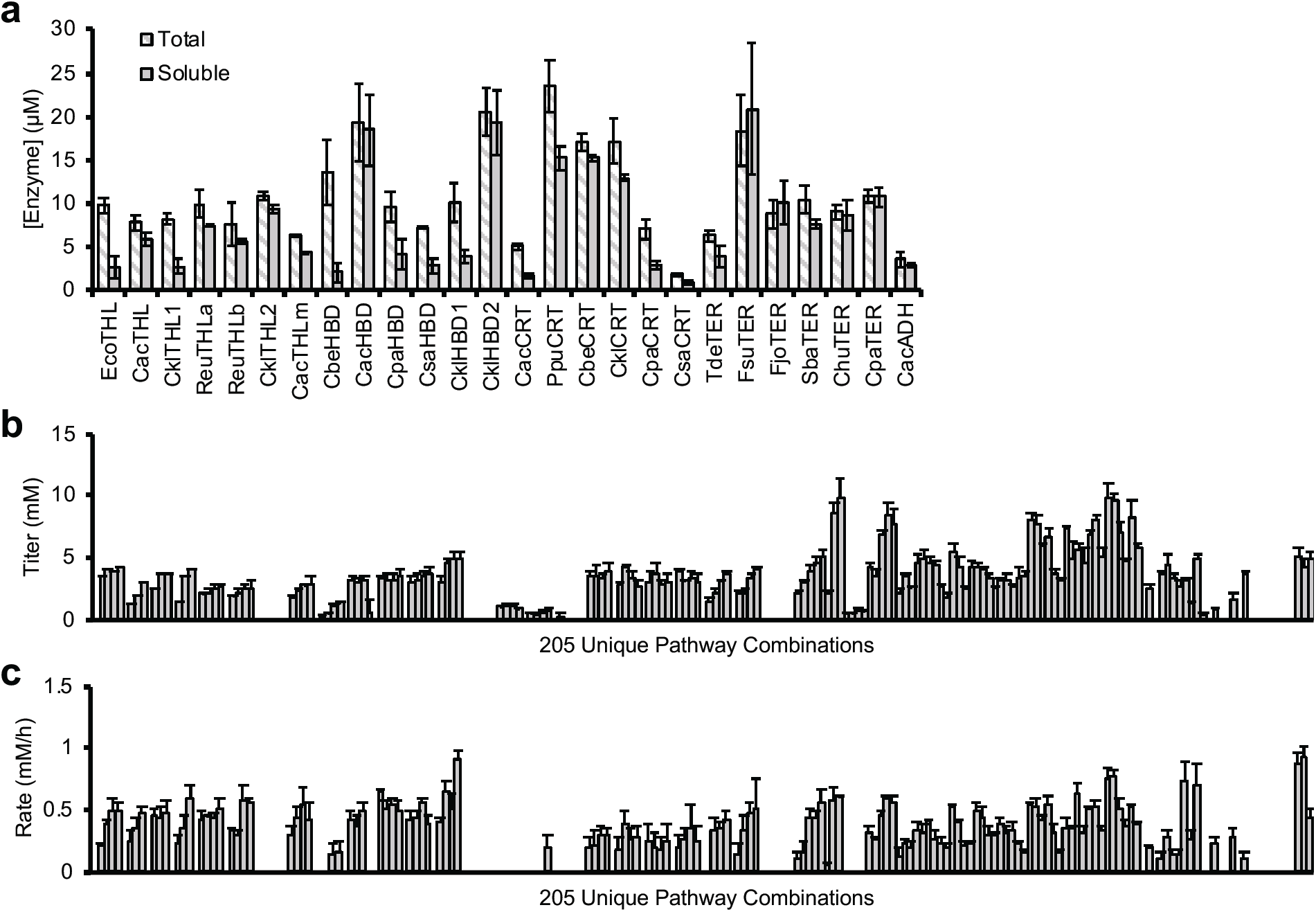
Cell-free butanol titers, rates, and enzyme expression. Six homologs have been selected for each reaction step (Thl, Hbd, Crt, and Ter) for *n*-butanol production and expressed by cell-free protein synthesis. Total and soluble yields of protein are displayed in panel (a) with error bars representing technical triplicates. iPROBE was run in 205 combinations listed in Supplemental File A. Titers at 24 h are shown in panel (b) with error bars representing technical triplicates. Rates determined by linear regression between 3 h and 6 h measurements are shown in panel (c) with error bars representing the standard error of the linear regression.

**Supplementary Figure S7.**
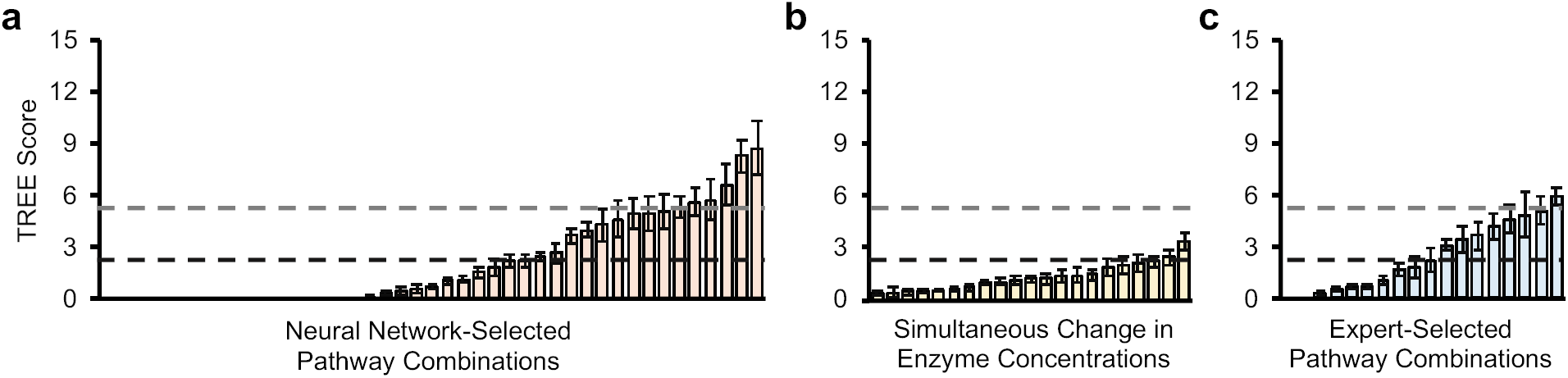
Cell-free experimental TREE scores for expert-selected and NN-based design of experiments. TREE scores were calculated for pathway combinations experimentally tested with iPROBE for (a) pathway combinations selected from the neural network approach (43 combinations), (b) simultaneous changes in each enzyme’s concentration using the base case set of enzymes (21 combinations), and (c) expert-selected pathway combinations based on data in Figure 5C and understanding of biosynthesis (18 combinations). TREE scores were calculated based on 24 h time-course data of *n*-butanol production.

**Supplementary Figure S8.**
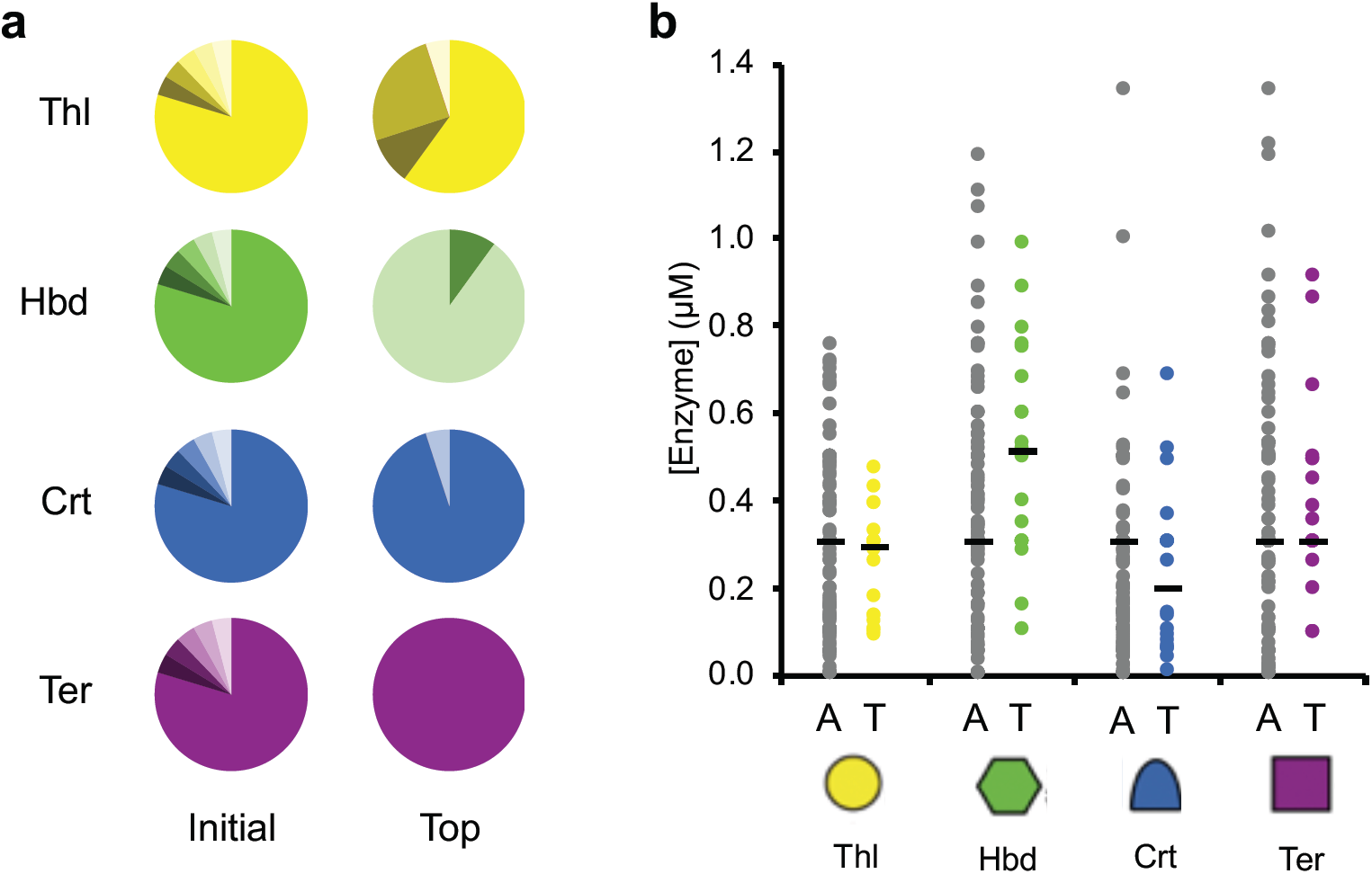
Analysis of iPROBE reaction compositions show enzyme homolog and concentration trends for butanol production. Each instance of each enzyme homolog of Thl, Hbd, Crt, and Ter tested in the 205 pathway combinations was counted. (A) The percentage of each homolog (six homologs total are represented for each enzymatic step) appearing in the ‘initial’ 120 combinations and in the ‘top’ 20 combinations (of all 205) is charted. (B) The concentrations used for each individual pathway enzyme (regardless of which homolog is used) in the assembly of all 205 combinations is plotted (A; grey) next to the concentrations used in assembling the top 20 combinations as ranked by TREE score (T; yellow for Thl, green for Hbd, blue for Crt, and purple for Ter). The median concentration for each enzyme and group is potted as a single black line.

**Supplementary Figure S9.**
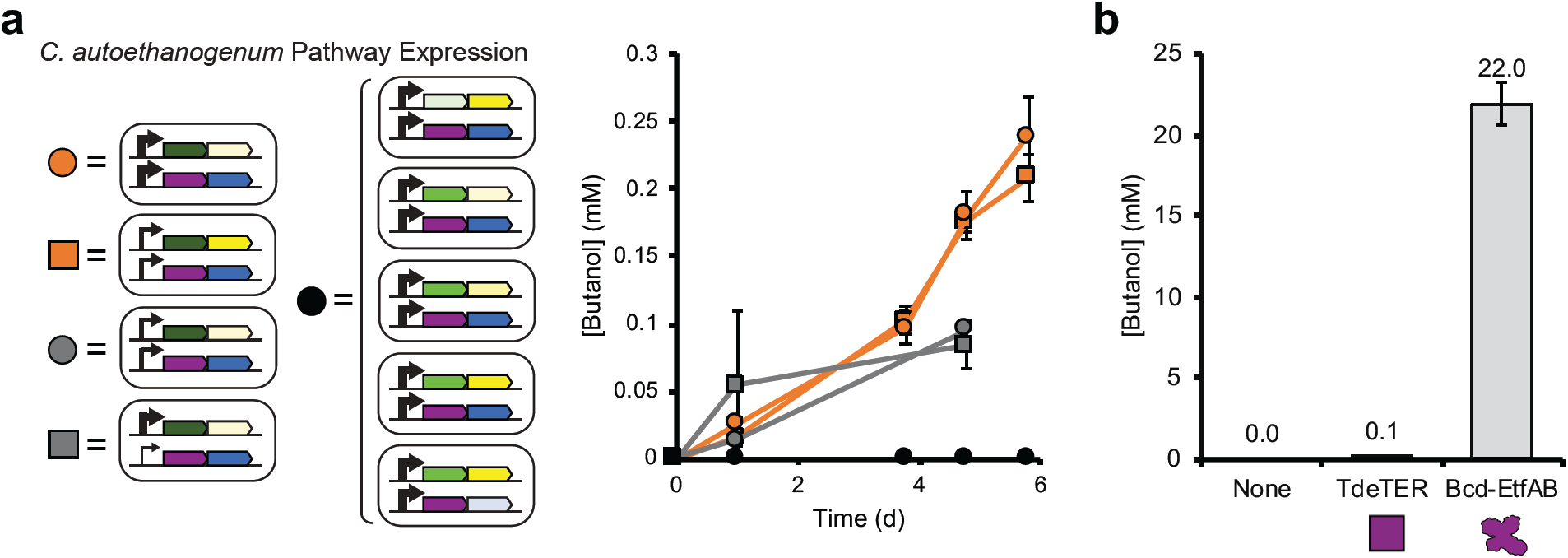
Cellular butanol expression in *C. autoethanogenum* is significantly enhanced when using Bcd:EtfAB versus *Tde*Ter. (a) Nine pathway combinations (two high-performing pathways, two medium-performing pathways, and five low-performing pathways) were constructed and transformed *in vivo* in *C. autoethanogenum*. Time-course measurements of butanol production were taken across six days and are plotted. Error bars for *in vivo* titers are standard deviations for four to six replicates. (b) Two pathways for *n*-butanol production containing *Cac*Thl, *Cac*Hbd, *Cac*Crt, and either *Tde*Ter or Bcd:EtfAB were built in high copy plasmids with the highest strength promoters in single operons. *C. autoethanogenum* strains containing these two pathway combinations were then fermented on gas and *n*-butanol was measured at 4 days. Error bars represent one standard deviation.

